# Multimodal evidence challenges the effectiveness of probabilistic cueing for establishing sensory expectations

**DOI:** 10.1101/2025.03.22.644773

**Authors:** Ziyue Hu, Dominic M. D. Tran, Reuben Rideaux

## Abstract

Predictive coding theories posit a reduction in error-signalling neural activity when incoming sensory input matches existing expectations – a phenomenon termed *expectation suppression*. However, the empirical evidence for expectation suppression, as well as its underlying neural mechanism, is contentious. A further aspect of predictive coding that remains untested is how predictions are integrated across sensorimotor domains. To investigate these two questions, we employed a novel cross-domain probabilistic cueing paradigm, where participants were presented with both visual and motor cues within a single trial. These cues manipulated the orientation and temporal expectancy of target stimuli with 75% validity. Participants completed a reproduction task where they rotated a bar to match the orientation of the target stimulus while their neural and pupil responses were respectively measured via electroencephalography and eye tracking. Our results showed a consistent, feature-unspecific effect of motor expectancy across multiple measures, while evidence for visual expectancy was limited. However, neither motor nor visual expectancy modulated the fidelity of sensory representations. These results indicate that violations of temporal expectancy in the current study may reveal the brain’s intrinsic sensitivity to temporal regularities in the natural settings, rather than feature-specific predictions. In contrast, the absence of visual expectancy effects in both neural and pupillometry results adds to a growing body of evidence questioning the effectiveness of probabilistic cueing paradigms for establishing expectations capable of altering sensory representations. Due to null findings in the visual and sensory representation analyses, we did not further investigate cross-domain prediction integration.

## 1. INTRODUCTION

Predictive coding frameworks challenge traditional views of perception as a bottom-up process, driven solely by the sensory reality (Kuffler, 1953). Instead, these frameworks propose that perception emerges through a dynamic interplay between incoming sensory inputs and pre-existing expectations (Friston, 2005, 2010; Lotter et al., 2020; Rao & Ballard, 1999). For example, our interpretation of an object’s convexity may depend on implicit assumptions regarding the typical direction of light sources (Sun & Perona, 1998). Within this framework, the brain is thought to continuously compare feedforward sensory input with internal generative models, whereby prediction errors – mismatches between input and expectations – propagate back through cortical hierarchies to update and refine these models (Friston, 2005). This iterative process is thought to optimize metabolic efficiency by minimizing the energy expenditure required for representing and engaging with the environment (Friston, 2005, 2010).

### 1.1. Expectation Suppression and Underlying Neural Mechanism

Predictive processing is substantiated by a growing body of evidence (de Lange et al., 2018; Kilteni et al., 2019; Kok et al., 2012, 2017; Rideaux et al., 2024; Summerfield et al., 2011; Tang et al., 2018a; Tran & Livesey, 2021). For instance, studies using functional magnetic resonance imaging (fMRI) and magnetoencephalography (MEG) show that auditory cues linked to specific visual stimuli can elicit anticipatory neural activity that closely resembles the patterns evoked by actual visual inputs, even in their absence (Kok et al., 2014, 2017). These findings highlight the brain’s active role in generating internal models to predict and interpret sensory information. The ability to form and leverage such expectations is thought to streamline processing demands (Friston, 2005, 2010), often evidenced by reduced neural activity in response to expected stimuli – a phenomenon known as *expectation suppression* (Kok et al., 2012; Summerfield et al., 2008; Tang et al., 2018). For example, transcranial magnetic stimulation (TMS) work in the motor domain reported reduced motor evoked potential (MEP) amplitude for predictable self-actions (Tran et al., 2021) relative to unexpected stimulation. Similar reduction in neural activity for expected events has been observed across other sensory modalities and neuroimaging techniques, including electroencephalography (EEG) measured event-related potentials (ERP) in auditory (e.g., N1, Harrison et al., 2021) and visual modalities (e.g., N1, Tang et al., 2018). However, the interpretation of expectation suppression effects requires careful consideration of potential confounds such as repetition suppression, a phenomenon characterized by similarly reduced neural responses but due to repeated presentation of identical stimuli (Grill-Spector et al., 2006). Cumulative evidence from the literature suggests that expectation and repetition suppression are at least partly distinct processes (e.g., Kaliukhovich & Vogels, 2011; Tang et al., 2018; Todorovic & de Lange, 2012). For example, Todorovic & de Lange (2012) demonstrated dissociable temporal profiles for expectation- and repetition-driven neural reductions. While such findings support a distinct role of expectation in shaping neural responses, the robustness of expectation suppression in the visual system has come under scrutiny by recent studies reporting null findings using probabilistic cueing designs (den Ouden et al., 2023, 2025). Such conflicting results indicate that predictive processing theories may benefit from considering alternative or complementary explanations, such as network fatigue models, which may more accurately account for some existing empirical findings (Feuerriegel et al., 2021; Feuerriegel, 2024). Therefore, despite the fact that predictive processing theories primarily attribute neural efficiency to top-down predictive mechanisms (Ali et al., 2022; Friston, 2005, 2010; Rao & Ballard, 1999), accumulating evidence suggests that other computationally-efficient low-level neural strategies may likewise contribute to the brain’s adaptive functioning (Mohan & Rideaux, 2025).

In addition to the debated robustness of expectation suppression, even in the literature where such effects have been observed, its underlying neural mechanism remains contested (Kok et al., 2012; Tang et al., 2018). One possible explanation is that the observed expectation suppression arises from the *sharpening* of neurons tuned to expected events, resulting in more precise neural representations compared to unexpected events (Kok et al., 2012; Kok & de Lange, 2015; Yon et al., 2018). Supporting this view, Kok et al. (2012) employed multivariate pattern analysis (MVPA) on fMRI blood-oxygen-level-dependent (BOLD) signals and found higher classification accuracy for expected (compared to unexpected) stimuli, suggesting that expected events are represented more precisely in the brain. This findings aligns with other neuroimaging evidence supporting the sharpening account (Den Ouden et al., 2008; Todorovic et al., 2011). Psychophysical evidence further corroborates this account; Kok et al. (2012) found that participants could more accurately detect differences in orientation and contrast when stimuli were expected. Similarly, Teufel et al., (2018) reported lower detection and discrimination thresholds for expected stimuli. These findings suggest that expected events are represented with better fidelity, consistent with the sharpening account of expectation suppression.

Although the sharpening account has gained some empirical support, an alternative hypothesis has been proposed in recent years; that is, expectation suppression may instead arise from the inhibition of neurons selective for expected features, leading to the reduction of representational fidelity for expected events (de Lange et al., 2018; Tang et al., 2018, 2023). This *dampening* of neurons tuned to the expected feature would instead result in unexpected events being represented relatively more precisely than expected events. Tang et al. (2018) investigated this account using inverted encoding modelling – a technique that predicts neural responses to specific stimulus features by mapping how these features are represented across continuous dimensions in the brain (Brouwer & Heeger, 2011; Harrison et al., 2023; Rideaux, 2024). Unlike MVPA, which simply attempts to discriminate between multiple neural patterns (Gessell et al., 2021; Kok et al., 2012), inverted encoding produces continuous feature-specific predictions, enabling finer analyses of sensory encoding. Using this approach, Tang et al. (2018) found higher representational fidelity for unexpected orientations compared to those that were expected, supporting the dampening account. Similar results have been observed across other studies on low-level features (Smout et al., 2019) and non-human animals (awake and anaesthetized mice, Tang et al., 2023). Therefore, contrary to the sharpening account of expectation suppression (Kok & de Lange, 2015), recent neuroimaging findings suggest that unexpected events may instead have better representational fidelity in the brain. Consistent with this hypothesis, Rideaux et al., (2025) used a task where participants replicated stimulus orientations by rotating a bar – allowing errors in these continuous responses to directly index representational fidelity – and found faster and more precise responses for unexpected stimuli. Similar behavioural evidence, corroborated by electrophysiological recordings, supporting the dampening account were found in another recent study investigating the influence of temporal context of visual processing (Lee & Rideaux, 2025). Thus, evidence from behavioural work seems to exhibit a similar inconsistency as that from neuroimaging experiments. However, it is important to note that the interpretation of multivariate decoding analyses, such as MVPA and inverted encoding modelling, depends on assumptions about the underlying neural populations (den Ouden et al., 2024) and their distinct temporal dynamics (Kok et al., 2014; Press et al., 2020). For example, decoding performance may reflect activity from prediction units, prediction error units, or a combination of both, each potentially exhibiting different temporal profiles related to feedback processing. In this study, and in much of the existing literature (Kok et al., 2012; Tang et al., 2018), representational fidelity is often interpreted as capturing prediction-modulated tuning of neuronal populations. Nonetheless, this interpretation remains largely implicit and has not been systematically tested; therefore, conclusions drawn from such results should be approached with appropriate caution.

### 1.2. Cross-domain Prediction

Understanding the nature of expectation suppression, a central theoretical and empirical tenet of predictive processing theories (Feuerriegel et al., 2021; Kok & de Lange, 2015), remains a challenging endeavor. Beyond its empirical robustness and competing accounts of its underlying neural mechanism, a yet unexplored question that may provide insight into predictive processing concerns how predictions from different sensory modalities (e.g., vision and audition) or domains (e.g., vision and motor) interact. Multimodal perception studies typically present participants with stimulus information from two different sensory modalities, and task them with responding to some aspect of the stimuli that is influenced by their integration. For example, Alais and Burr (2004) presented participants with both visual and auditory stimuli, manipulating the reliability of each to investigate how the brain integrates bimodal sensory information to infer the spatial location of a single event. While this form of multimodal experimental design has been used extensively to study how the brain integrates low-level features (Alais & Burr, 2004; Buhmann et al., 2024; Rideaux et al., 2021; Rideaux & Welchman, 2018; Stanford et al., 2005; Stein & Stanford, 2008), it has not yet been implemented within the context of predictive processing, namely, to study how the brain integrates higher-level sensory expectations. Several predictive coding studies have employed multiple sensory modalities or domains, but so far this has been applied in the context of considering predictions formed in one modality or domain which signals the event probability in another (Kok et al., 2012, 2014, 2017; Stekelenburg & Vroomen, 2015). For instance, Kok et al. (2012) used auditory cues to predict the orientation (45° or 135°) of visual target stimuli. Another study by Tran et al. (2025) simultaneously measured motor predictions using MEPs and auditory predictions using ERPs. They found that the strength of error signaling in one domain predicted the strength of error signaling in the other. The results suggest that predictions across different domains are related, and they support the notion that predictive coding may be governed by domain general mechanisms. However, this study did not investigate the fidelity of the neural representations or how predictions across domains were integrated. Thus, it remains unclear how predictions from different domains interact to represent a single event. In natural settings, the integration of prediction signals from multiple sensory modalities or domains is essential for a wide range of activities, from simple daily tasks (e.g., catching a falling object) to complex athletic performances (e.g., predicting the trajectory of a ball and timing a precise hit). These examples highlight the importance of adopting a cross-domain perspective on predictive processing.

### 1.3. Current Study

To address the ambiguous empirical landscape of expectation suppression and its underlying neural mechanism, as well as the gap in our understanding of cross domain predictions, this study employed a novel cross domain probabilistic cuing paradigm. We presented participants with both visual and motor cues, which respectively manipulated the featural (orientation) and temporal expectancy of the target stimuli with 75% validity. To assess the representational fidelity of (un)expected events, we combined inverted encoding modelling of EEG data with a continuous reproduction task to assess both neural and behavioural evidence, respectively. We hypothesized that participants’ neural decoding and behavioural responses would either be more precise for expected events, aligning with the sharpening account, or for unexpected events, based on the dampening account. In addition, we used a counterbalanced 2×2 (visual expected/unexpected, motor expected/unexpected) design to examine whether such expectancy effects from multiple domains combine additively, as observed for other forms of cue integration (Stanford et al., 2005).

Consistent with previous studies (e.g., Hall et al., 2018; Kok et al., 2012; Tang et al., 2018), we did not explicitly disclose the cue-target association to the participants. However, to cross-validate whether we had successfully manipulated expectation, we also tracked participants’ pupil diameter changes. Although pupil dilation can be modulated by a variety of cognitive and physiological factors (Alnæs et al., 2014; van der Wel & van Steenbergen, 2018) and is not specific to surprise, it has been consistently associated with the processing of unexpected events (Lavín et al., 2013), and has been shown to have predictive utility as a measure of surprise in previous studies (Mazancieux et al., 2023; Rideaux, Dang, et al., 2025). By simultaneously recording neural, behavioural, and physiological responses to expectation, we sought to provide a comprehensive view of how expectancy shapes sensory processing.

## 2. METHODS

### 2.1. Participants

58 first- and second-year psychology students were recruited from the University of Sydney, with five participants excluded for misunderstanding of the task and two for major technical corruptions (i.e., equipment disconnections between task and recording devices) of the EEG data. The final sample size (*n* = 51, M_*age*_ = 20.23, N_*male*_ = 7) was similar to previous experiments using similar neural decoding methods (Harrison et al., 2023; Rideaux, 2024; Rideaux et al., 2023). Since the experiment was largely visual, participants were required to have normal or corrected-to-normal vision (assessed using a standard Snellen eye chart). The study took approximately two hours and participants received two-unit credits for their participation. Participants were all naïve to the aims of the study, and the experiment protocol was approved by the University of Sydney’s Psychology Low Risk Human Ethics Committee.

### 2.2. Apparatus

The current study was conducted in a dark, sound-proof, and electromagnetically shielded room. Task stimuli were displayed on a mid-grey disc centred on a black background, to reduce orientation cues provided by the edges of the screen. Stimuli were presented on a display monitor (ASUS VG248QE 144 Hz 24’’ 1920×1080 HDMI Gaming Monitor) via MATLAB R2020a (The MathWorks, Inc., Matick, MA) and PsychToolbox (v3.0, http://psychtoolbox.org/; Pelli, 1997). Participants were instructed to place their head on a chin-forehead rest, controlling the viewing distance at 70 cm with the screen subtended to 41.47° x 23.33°.

### 2.3. Stimulus

Referring to **Figure 1**, the target stimulus consisted of two wedges arranged apex-to-apex, creating a bowtie configuration (orientation randomly sampled from between 0 and 180°), which focused participants’ attention on the screen centre. To establish visual expectations in participants, a visual cue was introduced before the target stimuli, consisting of two aligned white lines displayed outside the region of the target stimuli. This design was intended to prevent adaptation in the region of the visual field where the target was presented (Rideaux et al., 2023) and maintain participants’ focus on the centre of the screen. To encourage participants’ use of the visual cue, the visibility of the target stimulus was reduced by presenting it briefly (100 ms) at low contrast (0.1 relative to maximum screen contrast). Additionally, a motor cue – a green dot placed at the centre of the screen – was presented to prompt participants to press the left mouse button.

**Figure 1.**
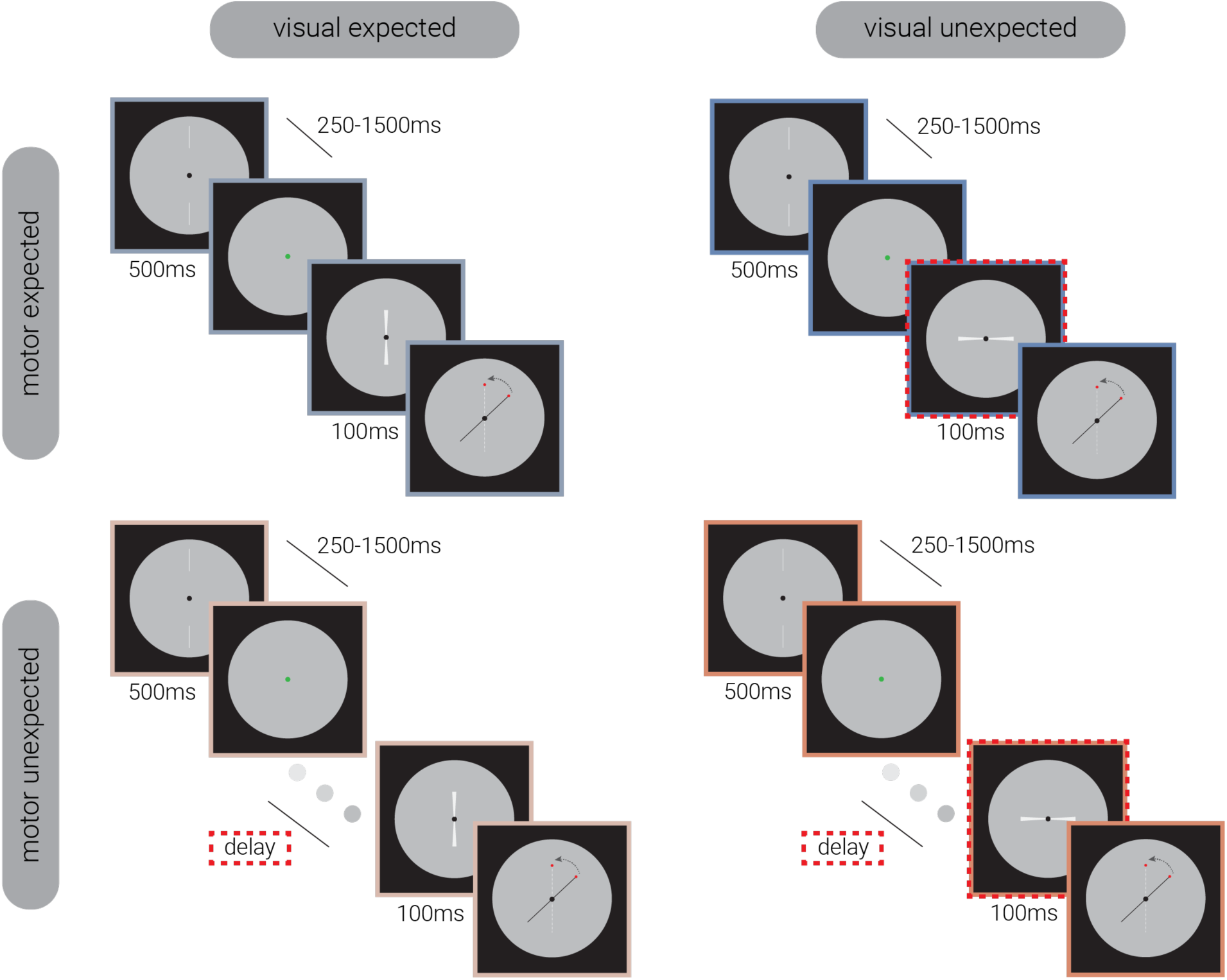
Experimental manipulation of visual and motor expectancy. Illustration of the trial design across conditions. Each trial began with a visual cue, followed by a variable delay (250-1500 ms) before a motor cue prompted participants to press the left mouse button. The target stimulus appeared either immediately (*motor expected; top row*) or after an additional variable delay (*motor unexpected; bottom row*), with its orientation either matching (*visual expected; left column*) or differing from the visual cue (>30°; *visual unexpected; right column*). Participants reported the target’s orientation by rotating a black line, with a red dot indicating the response index. The dashed red rectangles highlight the expectancy manipulations.

### 2.4. Design

Each trial began with the presentation of a visual cue for 500 ms, followed by a variable delay (250–1500 ms). Participants were then prompted by a motor cue to press the left mouse button. Upon release of the button, the target stimulus either appeared immediately (*motor expected, ME*) or after a variable delay (*motor unexpected*, *MU*) which was uniformly sampled from one-third of the motor-visual cue delay range. The orientation of the visual cue either matched the target (*visual expected, VE*) or differed significantly (>30°; *visual unexpected, VU*). Luminance was controlled across all target stimuli. The motor cue, however, differed in brightness from the subsequent target stimuli, producing a transient luminance change at target onset. Because visual expectancy effects were later analysed by averaging across motor expectancy conditions, the exposure to luminance change was balanced across trials and is unlikely to confound those analyses. In contrast, in the motor expectancy manipulation, the additional temporal delay meant that motor unexpected trials were exposed to the brighter cue for longer, which we note that could contribute to baseline pupil diameter differences between motor expectancy conditions. Participants reproduced the target’s orientation using a mouse-controlled black line, with the initial angular position randomized to prevent response bias. To facilitate responses, a red dot indicated the reproduction index during the response phase. There were no time limits for participants to complete the reproduction task.

There were 2 (visual expectancy: VE/VU) x 2 (motor expectancy: ME/MU) conditions in the current study (see **Figure 1**). Trials from each condition were pseudo-randomly interleaved within each block. There were 10 blocks in total, with 72 trials in each block. Visual expectancy trials comprised 75% expected trials and 25% unexpected trials. Similarly, motor expectancy trials contained 75% expected trials and 25% unexpected trials. The two conditions were counterbalanced. For example, 75% of visual expected trials were also motor expected trials (see **Figure 2**). This design allowed us to explore how sensory representations change dynamically when predictions from both visual and motor domains are met, only one of them is met, or both are violated (i.e., VEME, VEMU, VUME, VUMU).

**Figure 2.**
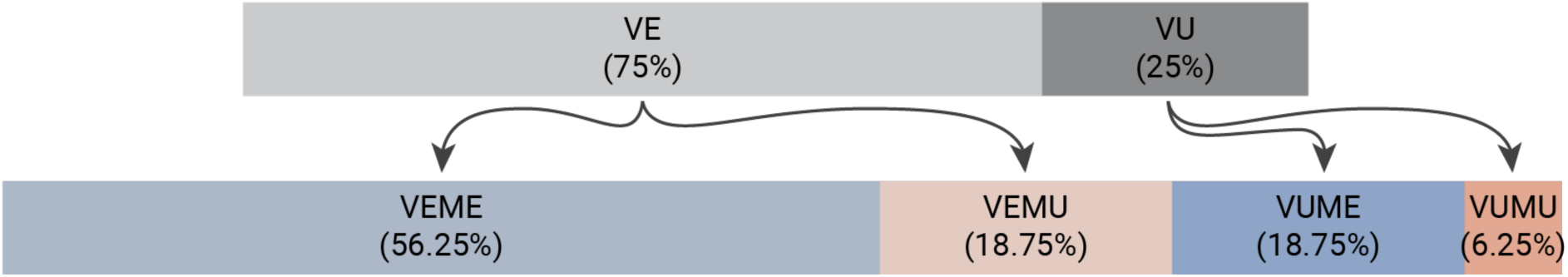
Illustration of counterbalancing four conditions (visual/motor expectancy). In each block, 75% of trials were in the visual expected condition, with these further divided into 75% motor expected and 25% motor violation trials. The remaining 25% of trials were visual violation trials, maintaining the same 3:1 distribution of motor expected and violation trials. Consequently, the visual expected/motor expected (VEME) trials constituted 56.25% of the total trials, while both the visual expected/motor unexpected (VEMU) and visual unexpected/motor expected (VUME) trials each made up 18.75%. The double-violation (VUMU) condition, involving both visual and motor unexpected trials, comprised 6.25% of the total trials.

There were two potential confounds that could have contributed to differences between expected and unexpected conditions. First, for each expectancy, there were three times more expected trials than unexpected trials, which could lead to higher EEG decoding accuracy in the expected condition. Second, the additional temporal delay between the visual cue and target in the motor unexpected trials may have, for example, disadvantaged behavioural reproduction of the target stimulus in these trials. These potential confounds were anticipated during the design of the experiment and to mitigate them, we used a variable visual-to-motor-cue delay (250 - 1500 ms), so that by omitting two thirds of trials from the expected condition with longer visual-to-motor-cue delays from the analysis, we could simultaneously match the number and duration of trials between conditions. That is, when performing two-way repeated measures ANOVA on the data, the number of trials was equated across the four conditions (∼45 trials per condition).

### 2.5. Procedure

Before the experiment, participants were provided with written and verbal instructions. They were informed that each trial would initiate with two aligned white lines, followed by the appearance of a green dot, prompting them to press the left mouse button. After their button-pressing response, the target pattern would appear, and the participant would be required to reproduce its orientation without time limits. Nevertheless, the relationships between visual/motor cue with the target stimulus were not explicitly disclosed. Participants completed a brief practice session with feedback (a red line indicating the correct orientation), but no feedback was provided during the testing session.

### 2.6. EEG Recording and Pre-processing

This study employed 64-channel active Ag/AgCl electrodes with the arrangement on the scalp aligning with the 10–20 international standard (Oostenveld & Praamstra, 2001), via a nylon cap and an online reference point at FCz. EEG data were captured at a 1024 Hz sampling rate using a 24-bit A/D conversion and recorded on a BrainVision ActiCap system (Brain Products GmbH, Gilching, Germany). The EEG data pre-processing was conducted using EEGLAB version 2021.1 (Delorme & Makeig, 2004). The data sampling rate was initially reduced to 256 Hz. To eliminate slow baseline fluctuations, a 0.1 Hz high-pass filter was applied, and a 45.0 Hz low-pass filter was employed to filter out high-frequency noise. Subsequently, independent component analysis (ICA) was performed using the automatic SASICA pipeline (Chaumon et al., 2015) to identify and remove components corresponding to ocular and other artifacts, including eye blinks. Subsequently, independent component analysis (ICA) was performed using the automatic SASICA pipeline (Chaumon et al., 2015) to identify and remove components corresponding to ocular and other artifacts, including eye blinks. Across the dataset, 21 posterior channels (1.6% of all channels) were excluded due to excessive noise and subsequently interpolated. A further 17 epochs (0.04% of the data) were lost due to recording errors (i.e., missed triggers). No additional data removals were performed. The final data were epoched from 200 ms to 500 ms relative to the onset of the target stimulus, and for post hoc analyses, the motor cue. The final data were epoched from −200 ms to 500 ms relative to the onset of the target stimulus, and for *post hoc* analyses, the motor cue.

### 2.7. Eye Tracking

Participants’ pupil responses were recorded via an Eyelink 1000 high-speed camera (version 4.594, SR Research Ltd., Mississauga, Ontario, Canada) that interfaced with an eye-tracking control monitor (Eyelink host PC). Participants’ pupillary responses during the experiment were monitored by the experimenter via a 1 kHz live video feed of the right eye displayed on the control monitor.

### 2.8. Data Analysis

#### 2.8.1. Behavioural Response and Mixture Modelling

In the behavioural reproduction task, we recorded both response time and replication performance (i.e., the angular difference between participants’ responses and the target orientation), with a focus on replication performance as the dependent variable of interest in the analysis. A preliminary analysis of circular standard deviation of replication performance (referred to as ‘*response error’*) revealed large standard errors in the current dataset. Therefore, to better understand the nature of errors, we utilized a mixture modelling approach similar to previous working memory literature (Zhang & Luck, 2008), to disambiguate trials on which participants were replicating the target orientation or merely guessing. Guess responses exhibit a uniform distribution of errors, showing no relationship with the target orientation. Conversely, valid replication of the target orientation would manifest as a distribution of errors centered on the target. The width of this distribution is inverse to participants’ behavioural precision, referred to as κ. A mixture model comprising a combination of these two distributions was fit to each participant’s data.

#### 2.8.2. Univariate Analysis of EEG data

Target stimuli in the current study were orientation-based, with orientation-selective neuron activities typically observed over occipital channels (Roe et al., 2012). Therefore, we defined an occipital region of interest (ROI) including all occipital and parietal electrodes.

#### 2.8.3. Inverted encoding Modelling

To identify the neural representations of the target stimuli, we utilized an inverted modelling approach to decode the target stimuli’s orientation from EEG recordings, consistent with previous work (Brouwer & Heeger, 2011; Harrison et al., 2023; Rideaux, 2024; Tang et al., 2018). We first developed a forward model that described the EEG activity in response to the target stimuli’s orientation. This model then served as a basis to build an inverse model that could interpret the EEG data to ascertain the orientation of the target stimuli. In the analysis, we applied a ten-fold cross-validation approach, wherein we trained the inverse model with 90% of the data set and then tested its accuracy by decoding the orientation from the remaining 10% of the data. In the current study, higher decoding accuracy was interpreted as indicative of more precise neural representations, allowing us to assess evidence for either the sharpening or dampening account. In the current study, higher decoding accuracy was interpreted as indicative of more precise neural representations, allowing us to assess evidence for either the sharpening or dampening account. Specifically, under the sharpening account, narrower neuronal tuning towards expected features should yield more precise representations and thus higher decoding accuracy for expected stimuli. By contrast, as the dampening account proposes reduced sensitivity to expected features, unexpected stimuli would instead be more distinctly represented and therefore associated with higher decoding accuracy.

Consistent with the previous literature (Harrison et al., 2023; Rideaux, 2024), the forward model consisted of five hypothetical channels with linearly spaced orientation preference between 0° and 180°. Each channel was expressed as a half-wave sinusoid raised to the fourth power. Therefore, the turning curve for each specific orientation would be described as a weighted sum of the five channels. The EEG activity observed for each presentation can be characterised using the following linear model:

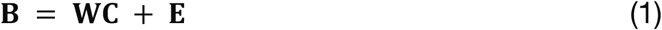

where **B** (*m* sensors x *n* representations) denotes the EEG data, **W** is a weight matrix (*m* sensors x 5 channels) to be estimated, **C** (5 channels x *n* representations) represents the nominated channel activities, **E** stands for residual errors.

To train an accurate inverse model, we sought to compute the weights that would reconstruct channel activities in EEG data with minimal error. Adapting from previous M/EEG work (Kok et al., 2017; Mostert et al., 2015), due to high correlations between signals recorded at adjacent sensors, we modified the conventional inverted encoding model approach to include noise covariance to enhance EEG data modelling accuracy. We also normalized components **B** and **C** to zero mean across presentations for each sensor and channel. The model was refined through a subset of data selected via the ten-fold cross-validation. Hypothetical responses for each of the five channels were derived from the training data, forming a response vector **c**_*train,i*_ for each channel ***i***, in proportion to the number of training presentations (**n**_*train*_). Sensor weights ***w***_***i***_ were then calculated using least squares estimation as following:

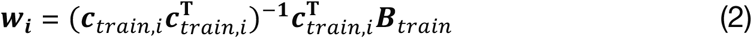

such that, ***B***_*train*_ (*m* sensors x *n* representations) denotes the EEG data training the forward model. Following this, we calculated the optimal spatial filter ***v***_***i***_ to reconstruct *i*th channel activity (Mostert et al., 2015) using:

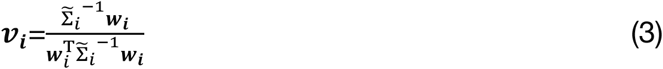

where 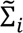 represents the regularized covariance matrix for channel *i*. Since adding noise covariance to the filter estimation could effectively reduce noise caused by high neighbouring sensor correlations, we estimated noise covariance in the following step. To enhance noise suppression, we further refined our estimation through regularization by shrinkage, which employed an analytically determined optimal shrinkage parameter (Mostert et al., 2015), forming a regularized covariance matrix 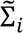 via:

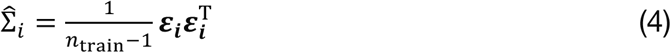

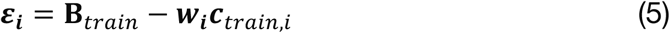

where **n**_train_ stands for the number of training representations. Adding up the spatial filters ***v***_***i***_ for all five orientation channels formed a channel filter matrix V (*m* sensors x 5 channels), which could then be used to reconstruct channel responses in testing set via:

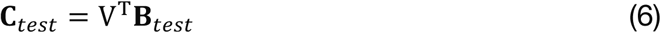

Subsequently, we decoded the orientation for each presentation by transforming the reconstructed channel responses to polar form as following:

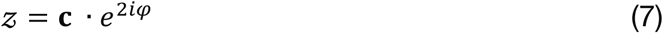

and then computing the estimated angle using:

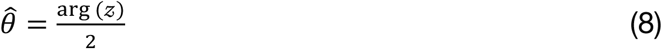

where ***c*** represents a vector of channel responses and ***φ*** is the vector of preferred angles for the channels, which was multiplied by two to map the 180° orientation space onto a 360° space. Based on the orientations decoded by our proposed model, we estimated the model’s decoding *accuracy*, defined as the similarity between the decoded orientation and actual presented orientation (Rideaux, Bays, et al., 2025). It was characterized by projecting the decoded-presented orientation differences onto a vector oriented at 0°:

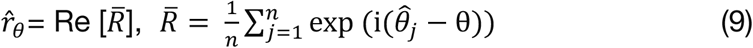

#### 2.8.4. Linear Discriminant Analysis

Linear discriminant analysis (LDA) is a dimensionality reduction technique used in classification to find a linear combination of features that optimally separates different classes (Subasi, 2020). After finding null effects in the inverted encoding modelling, we further performed a *post hoc* analysis, where we applied LDA to the EEG data to test whether the trials from the current dataset (visual and motor expectancy) could be classified based on trial expectancy (expected/unexpected). Since, for example, visual expected and visual unexpected trials each contained the same proportion of motor expected and motor unexpected trials (see **Figure 2**), we were able to average across the other expectancy dimension when testing classification for visual or motor expectancy. Classification accuracy indices were generated separately for visual and motor expectancy.

#### 2.8.5. Pupillometry data

Pupillometry recordings were epoched to a duration of −500 to 1500 ms around the target stimulus presentation. Data were pre-processed by removing blinks (100 ms buffer window), removing outliers (median absolute deviation>2.5), interpolated, bandpass filtered with a first-order Butterworth filter (highpass=15.6 Hz, lowpass=500 Hz), *z*-scored, epoched, and down-sampled to 125 Hz, respectively.

### 2.9. Statistical Analysis

#### 2.9.1. Behavioural Analysis

Repeated-measure ANOVAs (2×2: visual expectancy, motor expectancy) were conducted on response error, reproduction precision, and guess rate, with effect size calculated (Lakens, 2013). Bayes Factors (BF; Jeffreys, 1998) were also calculated to compare statistical support for both null and alternative hypotheses.

#### 2.9.2. Cluster Correction and Bayesian Repeated-measures ANOVA of Time-Series Data

EEG and pupillometry yield time-series data. Multiple comparisons at each time point can lead to a high risk of Type I error, so cluster-based permutation tests were employed to address this issue (Maris & Oostenveld, 2007). The analysis included 358 (time points) comparisons per test with 10,000 permutation samples, and a cluster-forming alpha of .05. For each permutation sample, four experimental conditions labels were shuffled and a 2 (visual expectancy) x 2 (motor expectancy) repeated-measures ANOVAs were conducted at each time point. Clusters were formed from *F*-values with *p*-values <.05, with adjacent time points treated as temporal neighbours. The sum of *F*-values within a cluster was defined as the cluster’s ‘mass’. The largest cluster masses from the 10,000 permutation samples formed the null distribution, against which the cluster masses from the original dataset were compared. The cluster-corrected significance of each cluster was determined by its percentile rank relative to the null distribution. This approach thus effectively controls the weak family-wise error rate while remaining sensitive to detecting effects in temporally autocorrelated EEG and pupillometry data (Groppe et al., 2011; Maris & Oostenveld, 2007). To test the effect of motor and visual expectancy on univariate and multivariate EEG data (inverted encoding modelling and LDA) and pupillometry responses, we conducted permutation repeated-measures ANOVA, and significant time periods were interpreted as those of mass exceeding the 95^th^ percentile of the null distribution. The Bayes factor for each time point was also calculated to evaluate evidence for both null and alternative hypotheses.

### 2.10. Data and code availability

The data and code for this study will be made publicly available on the Open Science Framework (https://osf.io) upon acceptance.

## 3. RESULTS

### 3.1. Effect of Visual and Motor Expectancy on Response Error

To investigate how expectancy modulates perceptual fidelity, we first analyzed the effects of visual and motor expectancy on response error (σ°), which was measured as the circular standard deviation of the angular differences between the target orientation and participants’ response. As shown in **Figure 3**, A 2×2 repeated-measures ANOVA revealed significant main effects of visual expectancy (*F*_1,51_=16.996, *P*<.001, *η*^2^ =0.250, *BF*_10_=245.351) and motor expectancy (*F*_1,51_=5.219, *P*=.027, *η*^2^ =0.093 *BF*_10_=1.747), with larger errors in unexpected trials, but no significant interaction effect (*F*_1,51_=0.364, *P*=.549, *BF*_10_=0.167), with Bayesian analysis providing substantial evidence favouring the null hypothesis.

**Figure 3.**
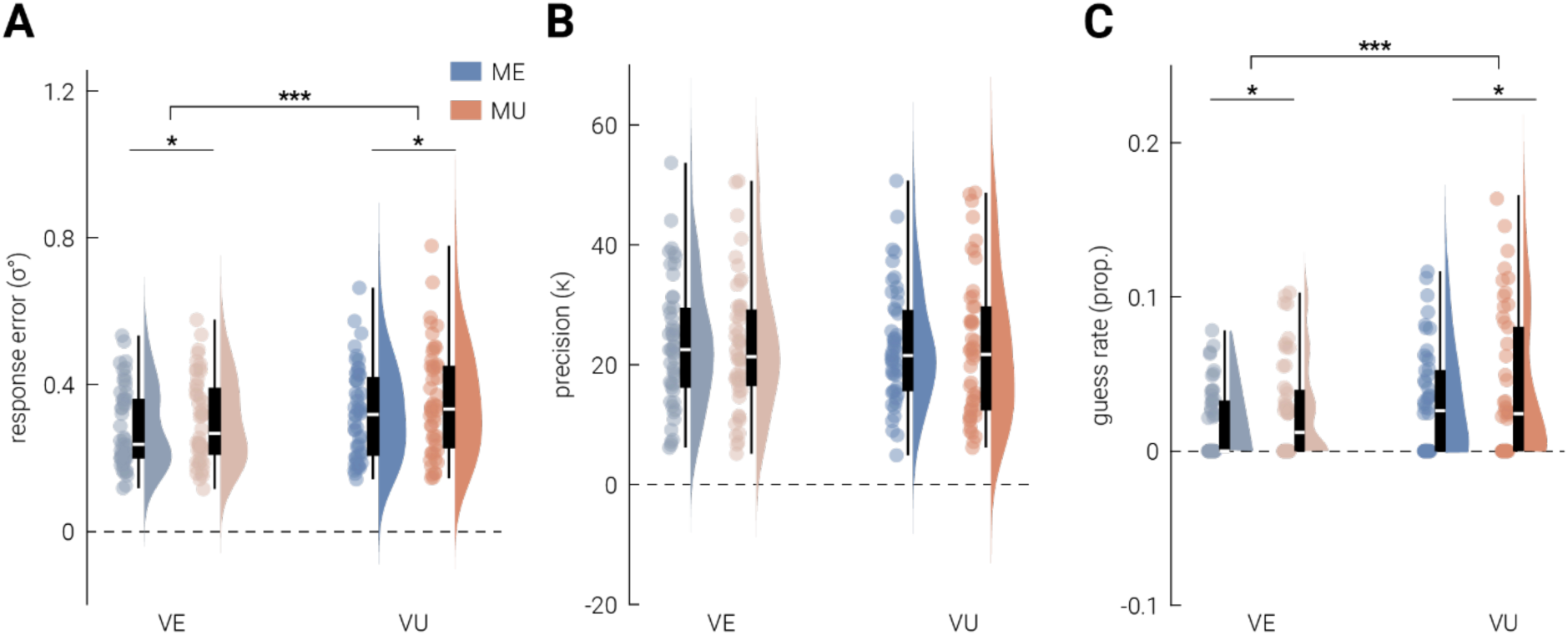
Behavioral response by visual and motor expectancy. **A**) Response error was calculated as the circular standard deviation of the angular differences between the target orientation and participants’ response. **B**) κ is an indicator of replication precision, which is inversely related to the width of the von Mises function fit to the data during mixture modelling. Higher κ value define narrower distributions, indicating more precise behavioral responses. **C**) Guess rate is defined as the frequency of participants’ guesses during the behavioral task, estimated from mixture modelling. Violin plots visualize the distribution of (**A**) response errors, (**B**) replication precision and (**C**) guess rate for each condition, with the median indicated by boxplots within each violin and semi-transparent dots indicating individual participant data points. Significant differences (p<.05 and p<.001) are marked with * and ***, respectively.

### 3.2. Response Error: Replication Precision and Guess Rate Analysis

To further investigate the source of the increased errors in violation trials, mixture modeling was used to differentiate participants’ behavioral responses based on replication precision and guessing (Bays & Husain, 2008; Zhang & Luck, 2008). **Figure 3B** illustrates the effects of visual and motor expectancy on replication precision (*κ*), where higher κ values indicate more precision. A 2×2 repeated-measures ANOVA revealed no significant main effect of visual (*F*_1,49_=0.250 *P*=.619, *BF*_10_=0.161) or motor expectancy (*F*_1,49_ =0.304, *P*=.584, *BF*_10_=0.165), both with substantial evidence for the null hypothesis indicated by the Bayes factor. There was also no significant interaction effect between two forms of expectancy (*F*_1,49_=0.031, *P*=.080) with the Bayes factor providing anecdotal evidence for the null hypothesis (*BF*_10_=0.859). However, for guess rate (frequency of guessing; **Figure 3C**), there were significant main effects of visual (*F*_1,49_ =12.930, *P*<.001, *η*^2^=0.209, *BF*_10_=49.340) and motor expectancy (*F*_1,46_ =4.295, *P*<.05, *η*^2^=0.081, *BF*_10_=1.156), with higher guess rates in unexpected trials than expected trials. The interaction effect was not significant (*F*_1,49_ =0.004, *P*=.950), with Bayesian analysis providing substantial evidence favoring the null (*BF*_10_=0.142).

### 3.3. Effect of Expectancy on Aggregated Neural Activity

Univariate analysis over the occipital-parietal ROI was conducted to investigate the effect of visual and motor expectancy on changes in the aggregated neural activity in this region. As shown in **Figure 4A**, there was no significant difference in neural responses driven by visual expectancy, but several time periods were observed for motor expectancy. In particular, there was a difference observed around 250 ms post-stimulus onset, exhibiting significantly greater P300-like activity in motor unexpected trials than expected trials. The latency and the scalp topography of this difference (inset in **Figure 4A**, characterized by peak amplitude mostly at central sensors) were consistent with typical findings of P3a, which is often linked to processing of novel events (Conroy & Polich, 2007; Polich, 2007). However, from - 200 ms to 100 ms around the target stimulus onset, neural responses in motor expected trials were significantly larger than those in motor unexpected trials. Given the timing of this difference, it could not be attributed to target stimulus processing.

**Figure 4.**
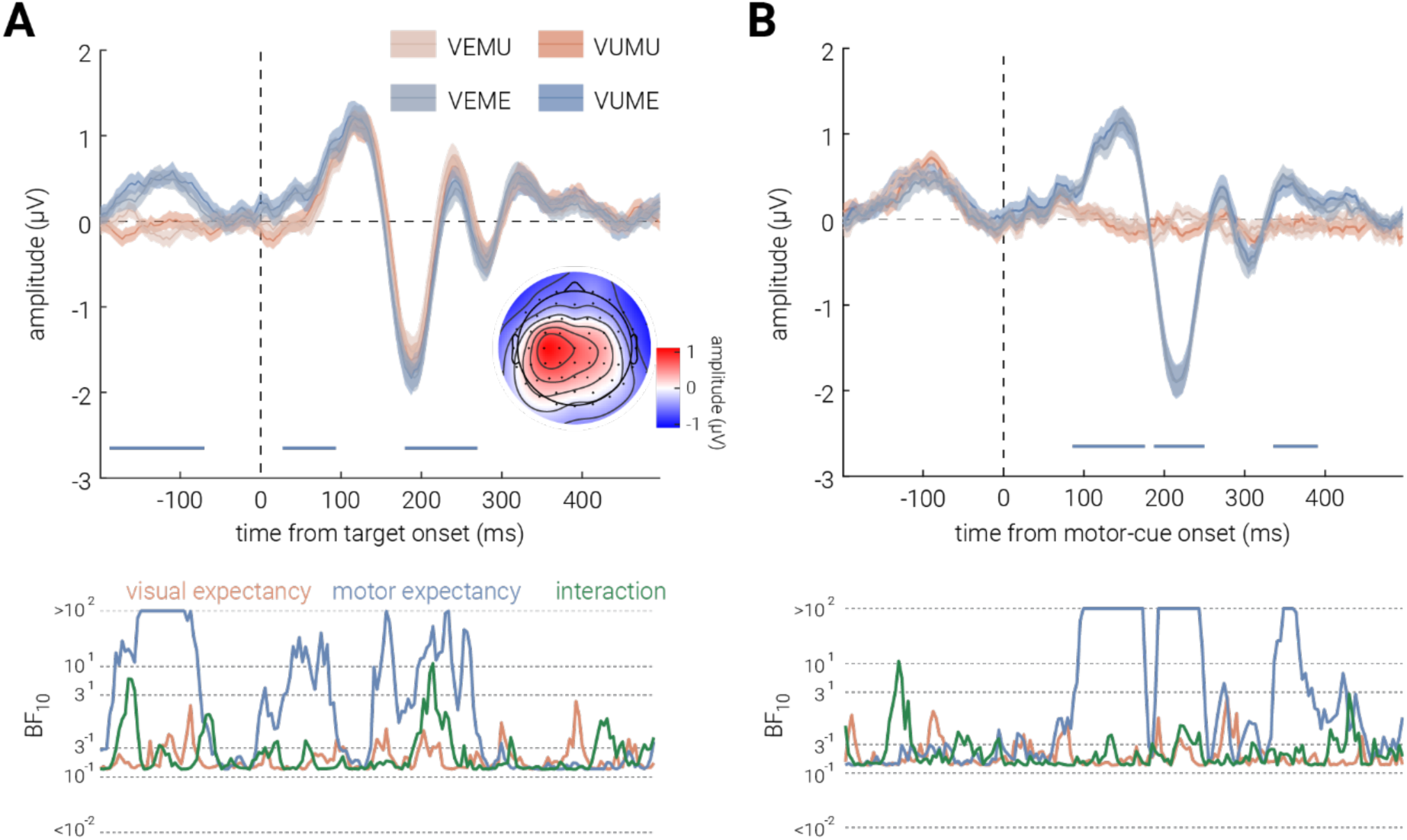
Effect of visual and motor expectancy on univariate EEG results. **A & B**) The effect of visual (VE, VU) and motor (ME, MU) expectancy on univariate EEG results when epoched to (**A**) target stimulus and (**B**) motor cue onset. The inset in (**A**) shows a topographic map of the average EEG activity across channels during the 180–280 ms post-stimulus interval, when a significant main effect of motor expectancy was observed. Shaded areas indicate ±1 SEM. Horizontal bars indicate cluster corrected periods of significant main effects of motor expectancy. The bottom sub-panels show the corresponding Bayes Factors (BF₁₀).

The univariate differences for motor expectancy in the first half of the epoch may be attributed to pre-stimulus baseline differences due to the temporal delay in target-stimulus onset introduced during motor unexpected trials. This pre-stimulus onset difference thus motivated us to conduct a *post hoc* analysis of neural activity epoched to 200 ms before and 500 ms after the onset of motor cue (**Figure 4B**). We found that the univariate neural response significantly differed by the motor expectancy after the motor-cue onset, specifically, a greater response was observed in the motor expected trials from ∼50 ms to ∼200 ms following the motor cue onset.

### 3.4. Effect of Expectancy on Neural Representation

#### 3.4.1 Stimulus Decoding: Inverted encoding Modelling

We found some evidence for a univariate motor expectancy effect, but none in the visual condition; however, univariate analyses can obscure meaningful multivariate changes in neural activity (Buhmann et al., 2024). Thus, to further explore whether expectancy influenced the neural representation of stimuli we employed inverted encoding to decode feature-specific neural representations, where higher decoding accuracy indicates more precise neural representation of the feature (orientation). As depicted in **Figure 5**, no significant differences in decoding accuracy were found for either visual or motor expectancy. The Bayes factors similarly provided anecdotal evidence for the null hypothesis, supporting the absence of an expectancy effect on stimulus representations.

**Figure 5.**
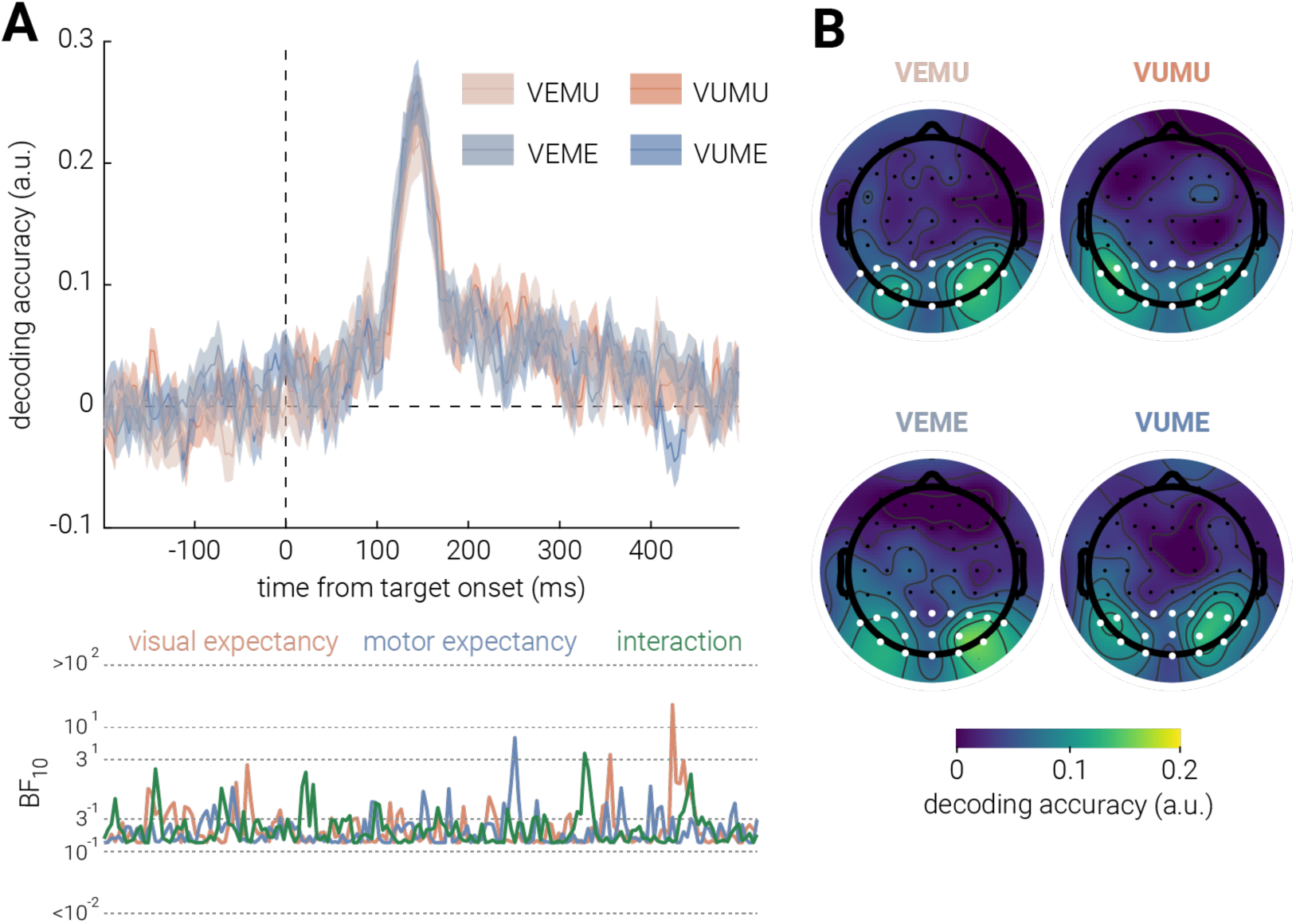
Effect of visual and motor expectancy on stimulus decoding. **A**) Decoding accuracy for the target stimulus orientation for visual (VE; VU) and motor (ME; MU) expectancy as a function of time from target onset. Shaded areas indicate ±1 SEM. The bottom sub-panels plot the corresponding Bayes Factors (BF_₁₀_). **B**) Topography of orientation decoding accuracy for the four conditions for the 0–400 ms interval following stimulus onset. The representation is largely clustered in occipital and parietal sensors. White dots indicate sensors included in ROI used for univariate and multivariate analyses.

#### 3.4.2 Expectancy Decoding: Post-Hoc Linear Discriminant Analysis

In our *a priori* analysis, we intended to examine the correlation between participants’ behavioral precision and feature-selective neural precision for target stimuli as a function of visual or/and motor expectancy. However, as no significant effects of expectancy were found in either behavioral or feature-specific neural precision, this analysis was not conducted. Instead, we performed a *post hoc* linear discriminant analysis on the EEG data to explore whether trial expectancy, rather than feature-specific representational fidelity, was encoded in the neural activity. That is, we tested whether there were feature unspecific differences in the multivariate pattern of activity between expected and unexpected trials, separately for visual and motor. As shown in the **Figure 6A**, there was no time periods when the classification accuracy of visual expectancy was significantly different from the chance level (50% accuracy). The Bayes factor provides similar anecdotal evidence supporting the null hypothesis. By contrast, classification accuracy for motor expectancy was consistently significantly above chance level for the entire epoch.

**Figure 6.**
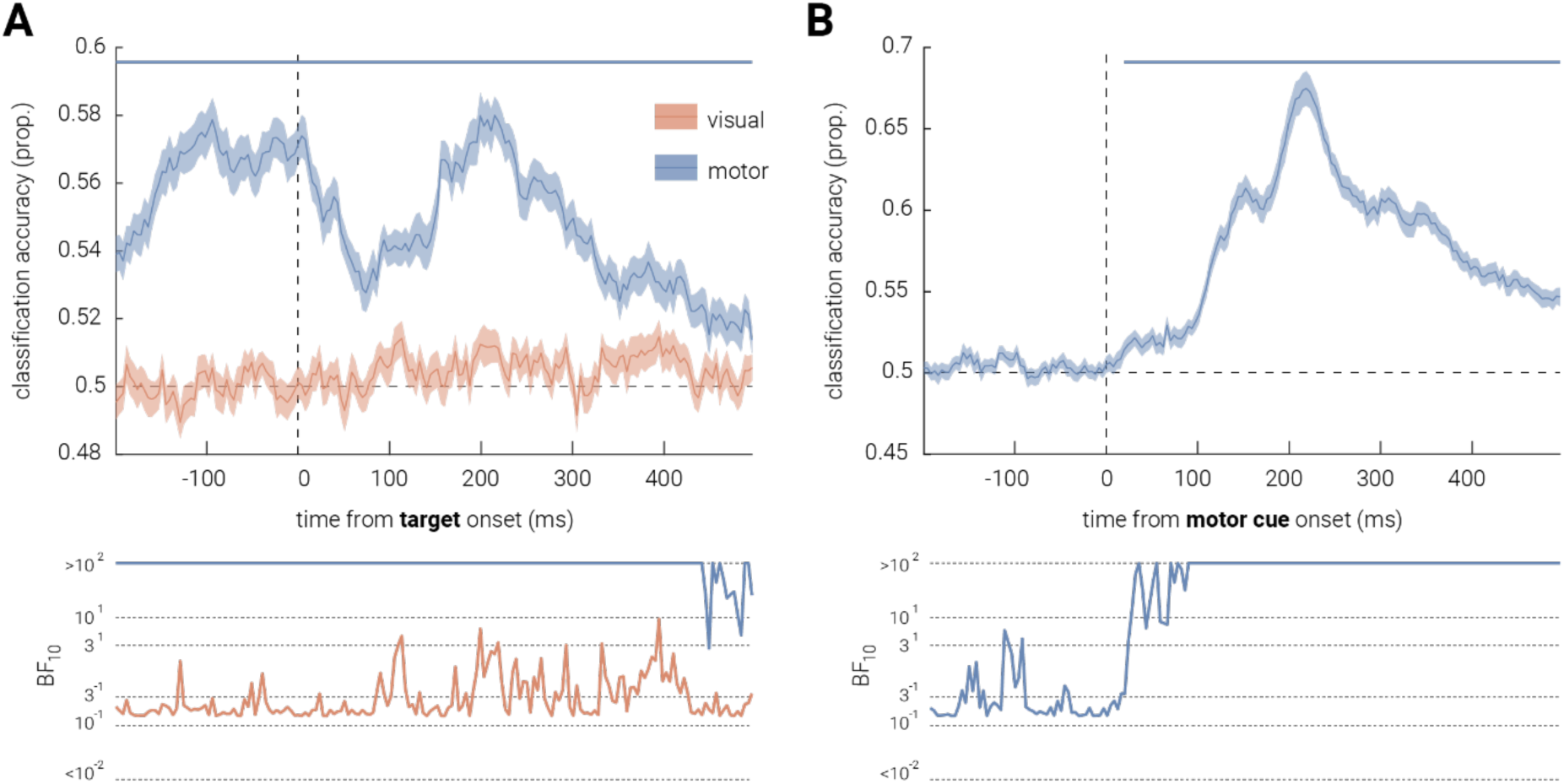
Expectancy decoding for visual and motor expectancy. Trial expectancy classification accuracy for (**A**) visual and motor expectancy, epoched to target stimulus onset, and for (**B**) motor expectancy only, epoched to motor cue onset. Shaded areas indicate ±1 SEM. Horizontal lines indicate cluster corrected periods of significant differences between classification accuracy and chance level (p < 0.05). The bottom sub-panel shows the corresponding Bayes Factors (BF₁₀).

The LDA results in the motor condition mirrored the significant pre-stimulus differences observed in the univariate EEG analysis. To further investigate the nature of these pre-stimulus differences, we re-epoched the EEG recordings to between 200 ms before and 500 ms after motor cue onset, aligning with the previous *post hoc* analysis. As illustrated in **Figure 6B**, classification accuracy for the motor condition was significantly above chance level, starting right after motor-cue onset with peak decoding accuracy at ∼200 ms post cue onset.

### 3.5. Effect of Expectancy on Pupil Response

In addition to neural and behavioural recordings, pupil diameter was recorded to cross-validate whether expectancy was successfully manipulated in our experimental design. We epoched the pupillometry recordings to between 500 ms before and 1500 after target-stimulus onset. Based on **Figure 7**, no significant difference in pupil diameter was observed for visual expectancy. The Bayes factor provides substantial evidence in support of the null hypothesis. However, pupil diameter was significantly larger for motor expected trials than unexpected trials from ∼200 ms following stimulus onset, with the Bayes factor suggesting similar decisive evidence for the alternative hypothesis. However, this pattern also appeared ∼500 ms before target stimulus onset and it was inverted, such that larger pupil diameter was found in motor unexpected trials than expected trials at ∼250 ms before stimulus onset. Notably, the increase in pupil diameter observed in motor unexpected trials, relative to motor expected trials, was significantly greater when the trial was also visual unexpected, with this interaction effect starting ∼600 ms after stimulus onset.

**Figure 7.**
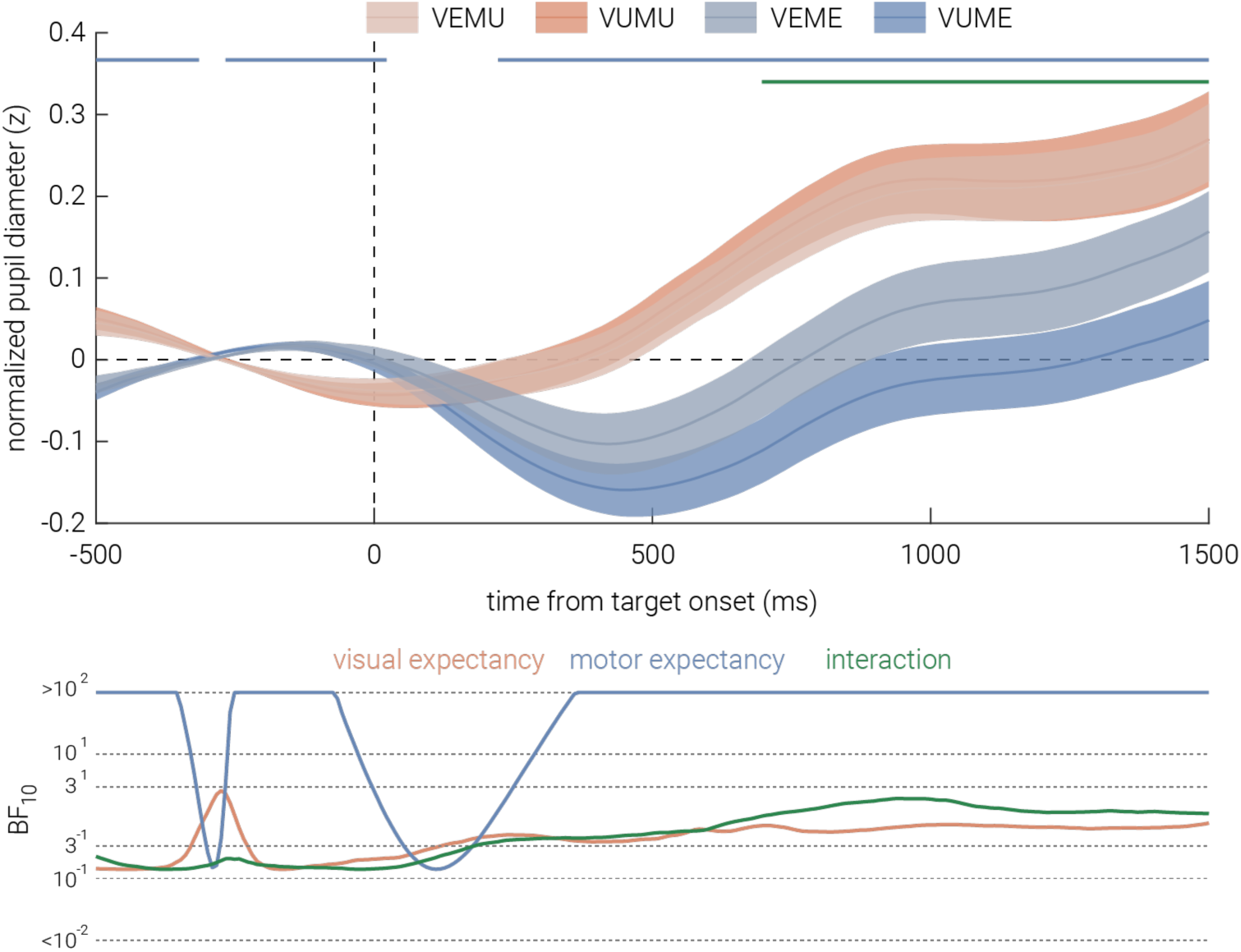
Effect of visual and motor expectancy on pupil diameter. Normalized pupil diameter in the visual (VE, VU) and motor (ME, MU) trials, as a function of time from target stimulus onset. Shaded areas indicate ±1 SEM. Horizontal bars indicate cluster corrected periods of significant main effects of motor expectancy and their interaction with visual expectancy (*p* < .05). The bottom sub-panels show the corresponding Bayes Factors (BF_₁₀_).

## 4. DISCUSSION

In this study, we aimed to test expectation suppression, its underlying neural mechanism, and how predictions interact across sensorimotor domains. Using a novel cross-domain probabilistic cueing paradigm, we combined EEG-based inverted encoding, a continuous reproduction task, and pupillometry, to assess neural, behavioral, and physiological responses to visual and motor expectancies. A seemingly persistent motor expectancy effect was observed across multiple measures, but it did not modulate sensory fidelity. In contrast, we found limited evidence for visual expectancy, aligning with recent null findings using similar probabilistic cueing paradigms (den Ouden et al., 2023, 2025). Given the lack of clear effects in both expectancies, the secondary aim of assessing cross-domain integration was not pursued. These findings assert implications for understanding the respective roles of intrinsic neural architecture and predictive processing in adaptive brain functioning, and inform directions for future experimental designs.

### 4.1. No Evidence for Expectancy-Driven Modulation on Sensory Representations

#### 4.1.1 Evaluating Expectation Suppression

Predictive coding theories propose that expectation suppression reflects reduced neural activity for expected stimuli, driven by decreased prediction-error signaling (Friston, 2005; Walsh et al., 2020). However, our univariate analyses did not reveal significant reductions in aggregated neural activity for visual expectancy. Similar findings have been reported in electrophysiological studies using probabilistic cueing paradigms for both high-level (e.g., faces, den Ouden et al., 2023) and low-level (Gabor gratings, den Ouden et al., 2024) visual stimuli. While the visual expectancy showed no evidence of expectation suppression, the motor expectancy exhibited a potential P3a suppression effect around 250 ms after stimulus onset (Polich, 2007). However, earlier neural activity (from 200 ms before to 100 ms after stimulus onset) showed the opposite trend. This reversal suggests that a pre-stimulus baseline difference may have influenced the later suppression effect. The pre-stimulus difference is likely due to the task’s temporal structure; inherent in our manipulation of motor expectancy is a delay to the onset of the target stimulus. In motor expected trials, the immediate onset of the target stimulus after button release likely carried residual motor and visual neural activity, whereas the delay in unexpected trials may have allowed this activity to stabilize. Regardless of the reason, the pre-stimulus difference makes interpretation of the post-stimulus difference difficult. Supporting this, a post-hoc univariate analysis re-epoched to the motor cue onset revealed larger neural responses in motor expected trials around 100 ms post-cue onset.

#### 4.1.2 Evaluating Neural Representational Fidelity

An alternative interpretation of the opposing pre- and post-stimulus trends is that the later P3a suppression may reflect a genuine expectancy-driven suppression superimposed upon a rebound artifact. However, univariate analyses may fail to capture subtle representational differences that could better explain these effects. Multivariate approaches offer a more sensitive method for examining how expectancy modulates sensory representations (Buhmann et al., 2024). However, our inverted encoding results revealed no significant differences in stimulus feature fidelity between expected and unexpected for both expectancies, suggesting that both types of events were encoded with similar precision in the current study, supporting neither sharpening nor dampening accounts. These findings appear to contradict hierarchical predictive coding models, which propose that prediction errors should modulate sensory representations (Friston, 2005; Kok & de Lange, 2015). Given the noticeable behavioural difference in response to expected and unexpected stimuli for both expectancies, the lack of decoding differences suggests that expectancy manipulations in the current study might not be effectively translated into neural predictions that alter sensory representations. However, recent efforts to consolidate the sharpening and dampening accounts suggest that both mechanisms may be in operation, potentially resulting in minimal net changes in neural fidelity. For example, the opposing processing theory (Press et al., 2020) proposes that the brain initially prioritizes expected stimuli, shifting toward unexpected stimuli only when strong expectancy violations occur. Thus, the null fidelity difference for both expectancies may reflect a co-existence of both sharpening and dampening of neurons selective to expected features. Nevertheless, if this were the case, given the sufficient temporal resolution of EEG, we would expect temporal transitions in decoding accuracy reflecting this shift, which were not observed in our results. Instead, current results are more likely explained by failure to elicit feature-specific neural expectations in the current probabilistic cueing paradigm.

#### 4.1.3 Behavioral and Physiological Expectancy Effects

While our inverted encoding modelling results provide no evidence for sharpening or dampening accounts, the behavioral and physiological data revealed feature-unspecific expectancy effects, particularly for motor expectancy. Mixture modelling of the reproduction task revealed that neither visual nor motor expectancy had effect on perceptual fidelity, aligning with neural analyses. However, unexpected trials elicited a higher guess rate for visual and motor expectancy, suggesting a general, feature-unspecific effect of expectancy on response behaviour. Rather than modulating sensory precision, unexpected events may disrupt attentional processes, leading to more frequent non-informative responses (Maunsell & Treue, 2006). This aligns with findings using attentional blink paradigms, where expectancy violations, particularly unexpected temporal delays, compromise task performance by interfering with attentional allocation (Tang et al., 2014).

Physiological data further reveals a similar motor expectancy effect. Pupil dilation responses were significantly larger in motor unexpected trials, consistent with prior research linking pupil size to surprise-related noradrenergic arousal (Lavín et al., 2013; Preuschoff et al., 2011). This result aligns with the behavioral and univariate neural findings, suggesting a potential motor expectancy effect in the motor condition that is not specific to a particular stimulus feature. However, these results must be interpreted with a note of caution. Significant pre-stimulus differences for motor expectancy were also observed; specifically at ∼250 ms pre-stimulus onset, there were larger pupil diameters in motor expected trials compared to unexpected trials. Given that pupil size inversely correlates with environmental luminance (Pan et al., 2022), the brightness contrast between the motor cue and target stimulus may have contributed to baseline diameter differences, potentially confounding later expectancy-related effects. This concern does not extend systematically to the analyses of visual expectancy effects, where exposure to the bright motor cue was balanced across conditions, but it should be acknowledged as a caveat when interpreting pupil effects related to motor expectancy. Nevertheless, despite pre-stimulus differences favouring larger pupil diameters in motor expected trials, we observed a greater post-stimulus increase in pupil diameter for motor unexpected trials compared to expected trials. Therefore, it remains plausible that the unexpected temporal delay in these trials elicited a surprise response beyond any baseline effects. Additionally, a significant interaction effect appeared around 600 ms after stimulus onset. Although this might initially appear to reflect visual unexpectancy enhancing the motor expectancy effect, closer examination showed that the interaction was primarily driven by a reduction in pupil diameter in visual unexpected trials rather than by increased dilation in motor unexpected trials. Given the late onset and direction of this effect, its functional significance is uncertain; however, it further suggests that visual expectancy was likely not established, as robust visual expectancy would be expected to elicit greater, not smaller, pupil dilation in unexpected trials. Consistent with this, no significant main effect of pupil dilation was observed for visual expectancy, mirroring the null results found in neural measures. Seemingly consistent neural, behavioral, and physiological findings in the motor condition tentatively suggest that motor expectancy may have elicited a feature-unspecific effect. In contrast, inconsistent evidence across the visual condition suggests that visual expectancy in the current study may not have been established.

### 4.2. Motor-Visual Discrepancy: Context-independent and Context-dependent predictions

One key pattern of the current results was that, across multiple measures, motor expectancy effects seemed to be consistent, as observed in increased guess rates and larger pupil dilation responses in motor unexpected trials, as well as a potential P3a suppression effect. However, null findings in behavioral precision or inverted encoding modelling suggests that these expectancy effects did not modulate the fidelity of sensory representations, contradicting both sharpening and dampening accounts (de Lange et al., 2018; Press et al., 2020). This suggests that the motor expectancy effect appeared to be feature-unspecific in the current study. Meanwhile, limited evidence for visual expectancy effects were observed. We further examined this motor-visual discrepancy via *post hoc* LDA, which revealed consistently above-chance classification accuracy for motor, but not visual, expectancy. Notably, pre-stimulus differences for motor expectancy mirrored univariate findings. To investigate this, we similarly re-epoched the motor LDA to the motor-cue onset, which exhibited significant above-chance classification accuracy shortly after the cue onset. Since LDA only captures differences between classes of data (Subasi, 2020), there are two possible interpretations for this post-cue result: on one hand, the difference could be driven by visual-field disparities between expected (seeing the target stimulus) and unexpected (not seeing until after a delay) trials; on the other hand, the difference might genuinely arise from the expectancy violation caused by the delay in unexpected trials (i.e., experiencing a delay when something is expected to occur immediately). These results reinforce the discrepancy between motor and visual expectancy in the current study: motor expectancy effects were possibly present albeit they didn’t modulate sensory representations, and the lack of robust LDA decoding in the visual condition further supports the absence of visual expectancy.

Feuerriegel (2024) argues that the brain’s adaptive functioning - especially in the visual system – can be primarily accounted for by low-level, spatiotemporal associations and the design principles of neural circuits, as proposed by network fatigue models. These models view adaptation as automatic, local mechanisms embedded in the structure of feature selective neurons and recurrent circuits, instead of top-down predictive processing or expectation-driven effects. Teufel and Fletcher (2020) similarly distinguish between context-independent and context-dependent predictions. The conceptualization of context-dependent predictions aligns with intuitive, high-level expectations, while context-independent predictions are defined as arising from the brain’s inherent architecture, reflecting statistical regularities in the naturalistic environment. Some may contend that the latter are not genuine ‘expectations’ in the traditional top-down sense (Feuerriegel, 2024), instead they better reflect automatic sensory adaptation based on one’s phylogenetic and ontogenetic development. However, this view can still be incorporated into predictive processing theories, especially hierarchical predictive coding models, which conceptualize feedback-based prediction as relative to the hierarchical brain organization, rather than from an absolute ‘high-level’ to ‘low-level’ (Kok & de Lange, 2015; Lotter et al., 2020; Rao & Ballard, 1999). Despite their taxonomical differences in prediction, both Feuerriegel (2024) and Teufel and Fletcher (2020) highlight that the brain’s circuit level design principles – shaped by natural environments – are as important, if not more so, for adaptive functioning as higher-level, context-dependent expectations.

In our study, unexpected temporal events induced by motor expectancy appear to support Teufel and Fletcher (2020)’s views on context-independent prediction. In particular, motor expectancy in the current study was based on temporal associations between an action (button press) and the appearance of a target stimulus. This form of prediction could be seen as context-independent, as such temporal regularities are stable across the daily environments and do not require explicit learning. The observed feature-unspecific motor expectancy effects may therefore reflect the brain’s intrinsic sensitivity to regular temporal associations rather than feature-specific predictions formed in the current paradigm.

However, the absence of neural effects for visual expectancy suggests two possibilities. First, in line with Feuerriegel (2024) and null expectation effects reported by recent studies (den Ouden et al., 2023, 2025), it might suggest that adaptive processes in the visual system, if observed, may indeed be better explained by local neural network mechanisms, rather than by higher-level expectation effects. Specifically, Feuerriegel (2024) argues that reductions in neural responses attributed to expectancy effects are often due to low-level sensory adaptation, such as stimulus-selective neural habituation. When such adaptation is controlled for, these reductions are typically absent. In our study, by spatially separating the visual cue from the target stimulus, we minimized the potential for low-level sensory adaptation. Consistent with Feuerriegel (2024), we did not observe significant neural response reductions for the visual expectancy manipulation, suggesting that previously reported effects may reflect uncontrolled sensory adaptation rather than genuine expectancy-based modulation. However, beyond previous work employing the probabilistic cueing paradigm, the current study incorporated an additional physiological measure to cross-validate expectancy manipulation. The absence of pupil dilation changes for visual expectancy suggests that, while the probabilistic cueing paradigm may reduce low-level adaptation confounds, its effectiveness in reliably establishing expectancy remains uncertain – potentially accounting for the null findings observed here and in prior studies. Therefore, the second possibility is that our probabilistic cueing manipulation may not have been sufficient to generate strong visual expectations. In this study, we manipulated the visual expectancy via cue-target associations across a continuous feature space (orientation). This form of prediction could be seen as context-dependent, requiring explicit reinforcement through repeated exposure to form strong associations. Supporting this, prior research has successfully induced context-dependent expectancy effects either using simpler, discrete cue-target mappings (e.g., two cue-target associations in Kok et al., 2012), or substantially larger trial numbers (e.g., 5400 trials in Tang et al., 2018). Thus, the lack of visual expectancy effects in this study may reflect insufficient exposure to establish robust context-dependent predictions. Future research could explore whether prolonged training, or restricted cue-target mappings, will assist a stronger establishment of context-dependent predictions. Taken together with our motor expectancy findings, these results contribute to the ongoing discussion regarding the extent to which the brain’s adaptive functioning is governed by intrinsic biological and neuronal architecture, and under what conditions higher-level, context-dependent predictive processes may (or may not) become engaged.

### 4.3. Conclusion

Predictive processing has long been investigated to understand how we navigate and adapt to the capricious environment with optimized efficiency (Friston, 2005, 2010; Kok & de Lange, 2015; Press et al., 2020). However, recent evidence and commentary have raised the possibility that previously reported expectation suppression effects may, at least in part, reflect low-level sensory adaptation, as these effects are often ambiguous when experimental designs sufficiently control for such confounds (den Ouden et al., 2023, 2025; Feuerriegel, 2024; Feuerriegel et al., 2021). Our findings show that violations of temporal expectancy in the motor condition potentially elicited feature-unspecific effects across multimodal measures, but they did not alter sensory representations. This pattern may reflect the brain’s context-independent sensitivity to temporal regularities commonly encountered in naturalistic settings, rather than higher-level expectations. In contrast, for higher-level, context-dependent visual expectancy, we found limited evidence, in line with recent work reporting null expectancy effects using similar probabilistic cueing paradigms (den Ouden et al., 2023, 2025). In our study, the experimental design minimized the influence of low-level sensory adaptation, supporting the interpretation that adaptive neural processing previously attributed to visual expectancy may actually be better explained by local neural mechanisms. However, our null pupillometry results offer an alternative explanation to the existing literature: the probabilistic cueing paradigm itself may not have been sufficient to reliably establish visual expectancy in the first place. Collectively, these results further inform ongoing debates regarding the relative contributions of intrinsic neural architecture and higher-level, context-dependent expectations in the brain’s adaptive functioning.

## Acknowledgements

This work was supported by an Australian Research Council Discovery Early Career Researcher Award and (DE210100790) a National Health and Medical Research Council Investigator Grant (2026318) to RR, as well as an Australian Research Council Discovery Early Career Researcher Award (DE220100829) to DT.

